# Long-term dynamic profiles of cognitive kinases induced by different learning protocols

**DOI:** 10.1101/2025.10.12.681952

**Authors:** Yili Zhang, Rong-Yu Liu, Paul Smolen, John H. Byrne

**Affiliations:** Department of Neurobiology and Anatomy, W.M. Keck Center for the Neurobiology of Learning and Memory, McGovern Medical School at the University of Texas Health Science Center at Houston, Houston, TX 77030

## Abstract

Learning is associated with activation of multiple protein kinases, but few details are known about the activation dynamics in response to different learning protocols. We addressed this issue by examining the long-term dynamics of kinases critical for long term synaptic facilitation (LTF) of the *Aplysia* sensorimotor synapse. Three serotonin (5-HT) protocols have been found to induce LTF with distinct effectiveness: the five-pulse regular-spaced Standard protocol; the five-pulse irregular-spaced Enhanced protocol; and the two-pulse protocol with an interval of 45 min. We previously compared long-term dynamics of the mitogen-activated protein kinase (MAPK) isoform ERK after these protocols. Here we examined the long-term dynamics of additional kinases critical for LTF: p38 MAPK, protein kinase A (PKA), and p90 ribosomal S6 kinase (RSK). All four kinases showed complex dynamics of activity during 24 h, with a first wave of increase occurring shortly after 5-HT treatment and ending within 5 h, and a second wave from ∼ 5 h to 18 h. After the Standard and two-pulse protocols, all kinase activities returned towards basal at 24 h, but after the Enhanced protocol some remained elevated at 24 h. Interactions and multiple feedback loops among the kinase pathways, and with the growth factors *Aplysia* neurotrophin (NT) and transforming growth factor-β (TGF-β), contribute to development of molecular clocks underlying these complex dynamics. These results help to delineate the molecular mechanisms underlying the induction of LTF and provide insights that may help design improved training protocols for induction and maintenance of LTF and long-term memory.

## Introduction

Learning leads to the activation of multiple protein kinases (Purcell et al. 2003; Ajay and Bhalla 2007; Pagani et al. 2009; Zhang et al. 2011, 2017, 2021; Liu et al. 2013, 2020; Jain and Bhalla 2014; Wang et al. 2014; Kopec et al. 2015; Mirisis et al. 2016; Kukushkin and Carew 2017; Zobon et al. 2021). However, the dynamics of kinase cascades critical for the formation of long-term memory (LTM) have not been rigorously quantified. Long-term synaptic facilitation (LTF) of the *Aplysia* sensorimotor synapse is a well-established system for studying the mechanisms underlying LTM (reviewed in Byrne and Hawkins 2015). Three serotonin (5-HT) protocols induce LTF: the Standard protocol of five, 5-min pulses of 5-HT with regular interstimulus intervals (ISIs) of 20 min; the Enhanced protocol of 5 pulses of 5-HT with computationally designed irregular ISIs; and the two-pulse protocol of two 5-min pulses of 5-HT with an ISI of 45 min (Zhang et al. 2011, 2023). The Standard and two-pulse protocols produce comparable LTF at 24 h, whereas the Enhanced protocol produces stronger LTF at 24 h and also prolongs LTF up to 5 days.

Essential roles in LTM, and associated synaptic plasticity, are fulfilled by multiple kinases, which are common to many cellular processes. Those that appear particularly critical for learning and memory have been called “cognitive kinases” (Schwartz, 1993). In *Aplysia*, PKA and ERK activated by 5-HT converge to regulate the transcription activator CREB1 and repressor CREB2, which in turn regulate genes (e.g., *c/ebp*) critical for LTF (Alberini and Kandel 2014; Byrne and Hawkins 2015); whereas p38 MAPK suppresses LTF by activation of CREB2, and its activity is biphasically regulated by 5-HT (Guan et al. 2002, 2003; Fioravante et al. 2004; Zhang et al. 2017). p90 ribosomal S6 kinase (RSK) is activated by the PKA and ERK pathways and contributes to CREB1 activation and LTF (Liu et al. 2019; Zhang et al. 2021). Positive feedback loops that include the growth factors *Aplysia* neurotrophin (NT)/Trk receptor and Tolloid/BMP-1/transforming growth factor-β (TGF-β) are recruited during learning to regulate the activities of PKA and MAPK (Zhang et al. 1997; Kopec et al. 2015; Jin et al. 2018). These kinases also are important for the formation of memory in other species, and reduction of these kinase activities leads to cognitive deficits (e.g., Goosens et al. 2000; Schafe and LeDoux 2000; Schafe et al. 2000; Delaunoy et al. 2006; Pagani et al. 2009; Rawashdeh et al. 2016). Our previous work demonstrated that optimizing the overlap of activities of PKA and ERK not only enhances LTF and LTM in *Aplysia* (Zhang et al. 2011) and but also enhances the acquisition of fear learning and extinction in rats (Zhang et al. 2023b).

Zhang et al. (2023) characterized the long-term dynamics of ERK activity after the three LTF-inducing protocols. A striking finding was a second wave of active phosphorylated ERK (pERK) that appeared to peak at 18 h post training. Also striking was that whereas pERK declined back to baseline at 24 h for the Standard and two-pulse protocols, it remained elevated at 24 h after the Enhanced protocol. These findings raised several key questions. First, do the dynamics of other key kinases such as PKA, RSK, and p38 MAPK show similar patterns of activity? Will they also remain activated at 24 h only after the Enhanced protocol? Second, what are the interactions between molecular pathways during the second wave of increase? Third, what are the role of the growth factors *Aplysia* NT and TGF-β in the second wave of increase? To address these questions, we characterized the dynamics of kinase activity up to 24 h after these protocols, and applied inhibitors to illuminate interactions between kinase pathways and growth factors.

## Results

### I. Dynamics of Kinase Cascades

#### 1. LTF-inducing 5-HT protocols induce two waves of increase in p-p38 MAPK within 24 h after 5-HT treatment

p38 MAPK functions as an inhibitory constraint for the induction of LTF (Guan et al. 2003; Zhang et al. 2021; Liu et al. 2022). To investigate the dynamic profile of p38 MAPK activity, we compared with time-matched vehicle controls, 5-HT-induced changes in the levels of active, phosphorylated p38 MAPK (p-p38 MAPK) immediately (0 h), 1 h, 2 h, 5 h, 18 h, and 24 h after the three protocols. These same time points were used to measure pERK in Zhang et al. (2023) and are used below to measure PKA and RSK activity, thereby allowing comparison of the dynamics of all four kinases. The Standard (Fig. 1A, grey line), Enhanced (Fig. 1A, red line) and two-pulse (Fig. 1A, blue line) protocols all induced two waves of increase in p-p38 MAPK. At the end of the 85-min Standard protocol (here defined as time = 0), p-p38 MAPK was increased to 44.8 ± 9.6% of vehicle control. At 1 h post treatment it was increased to 27.9 ± 8.3% of control, but declined at 2 h (−9.8 ± 8.3%) after treatment. p-p38 MAPK increased again at 5 h (23.8 ± 7.4%), the increase was also detected at 18 h (34.6 ± 6.9%), followed by a return to basal level at 24 h (3.6 ± 6.4%). Detailed statistics for experiments for which significant effects were found are given in Table 1.

**Figure 1.**
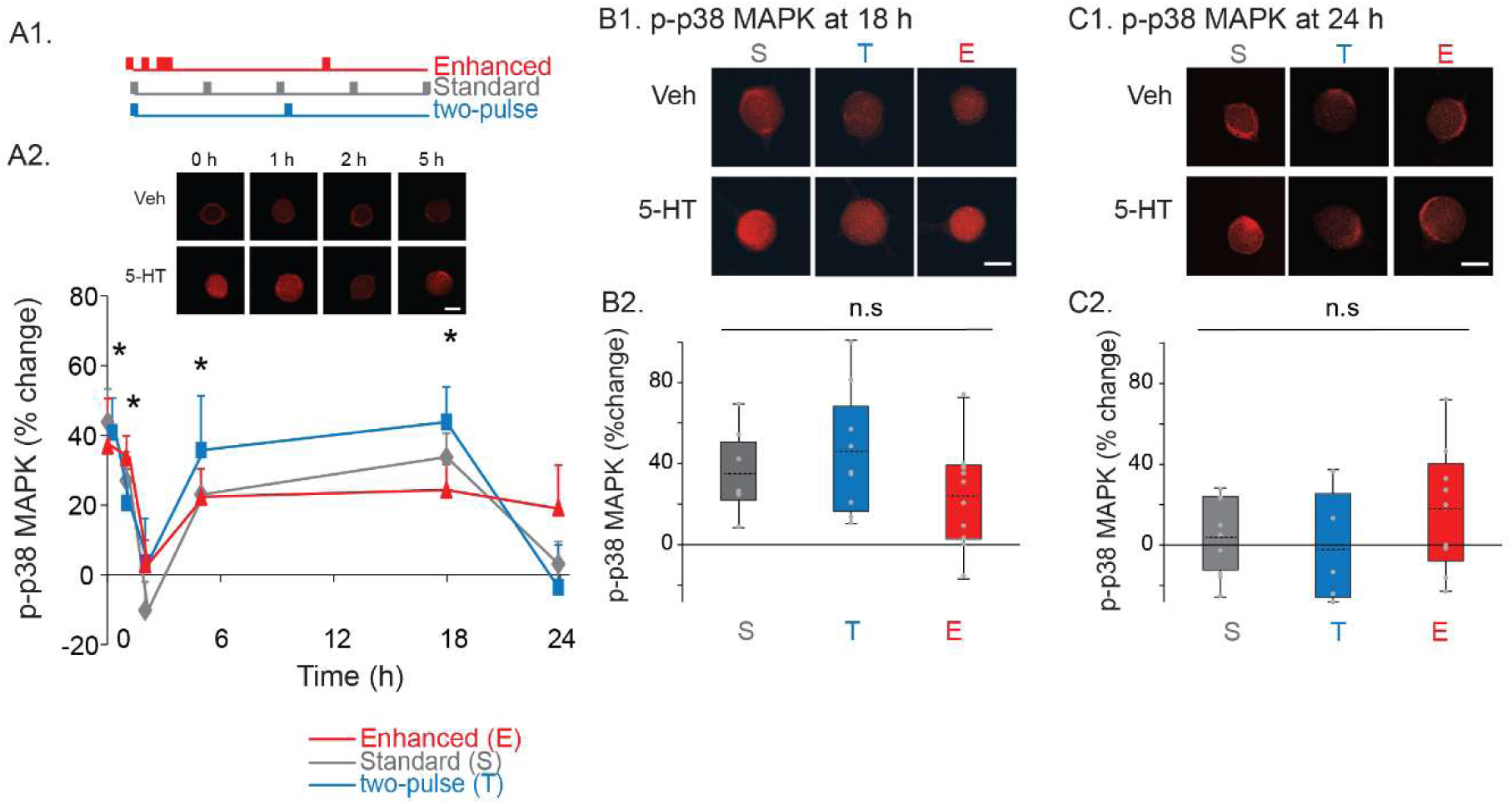
Dynamics of p-p38 MAPK induced by three LTF-inducing protocols. ***A1***, three LTF-inducing protocols. ***A2***, Time course of p38 MAPK activity, with representative confocal images (top) and summary data (bottom). The percent change was calculated as the change of p-p38 MAPK level after 5-HT compared to time-matched vehicle control. For all protocols, p-p38 MAPK was increased immediately (0 h) after 5-HT and at 1 h after, returned to near basal levels at 2 h, and then increased again at 5 h. This increase was still evident at 18 h for all protocols, followed by a return towards basal level at 24 h. p value was calculated by t-test analysis of 5-HT treated vs. Veh groups at each time point. * p < 0.05. ***B***, Comparison of p-p38 MAPK at 18 h after different 5-HT protocols (data from panel A). ***B1***, Representative confocal images of p-p38 MAPK immunostaining in SNs. In this figure and subsequent figures, all scale bars are 40 μm. ***B2***, Summary data. Statistical analyses did not reveal significant differences between p-p38 MAPK at 18 h induced by three protocols. ‘n.s’ represents ‘not significant’. ***C***, Comparison of p-p38 MAPK at 24 h after different 5-HT protocols. ***C1***, Representative confocal images of p-p38 MAPK immunostaining in SNs. ***C2***, Summary data. Statistical analyses did not reveal significant differences between p-p38 MAPK at 24 h induced by three protocols. Here and subsequently, the summary data are presented by box-and-whisker plots. The mean is indicated by the dashed line in the interior of the box. The lower end of the box is the first quartile (Q1). The upper end of the box is the third quartile (Q3). Circles indicate the results of individual experiments.

**Table 1.** Statistical analyses of Figures 1-8.

| **Fig. 1A, Standard protocol, paired t-test, Bonferroni correction** |  |  |  |
| --- | --- | --- | --- |
|  | t | p | n |
| 0 h | -6.276 | <0.01 | 10 |
| 1 h | -3.822 | 0.018 | 11 |
| 2 h | 1.619 | 0.864 | 9 |
| 5 h | -3.148 | 0.048 | 14 |
| 18 h | -4.831 | 0.012 | 8 |
| 24 h | -0.622 | 1 | 9 |
| **Fig. 1A, Enhanced protocol, Wilcoxon Signed Rank Test, Bonferroni correction** |  |  |  |
|  | z | p | n |
| 0 h | 2.521 | 0.048 | 8 |
| 1 h | 2.521 | 0.048 | 8 |
| 2 h | -0.280 | 1 | 8 |
| 5 h | 2.521 | 0.048 | 8 |
| 18 h | 2.578 | 0.042 | 11 |
| 24 h | 1.007 | 1 | 9 |
| **Fig. 1A, two-pulse protocol, paired t-test, Bonferroni correction** |  |  |  |
|  | t | p | n |
| 0 h | -4.162 | 0.036 | 7 |
| 1 h | -3.973 | 0.042 | 7 |
| 2 h | 0.188 | 1 | 8 |
| 5 h | -3.520 | 0.048 | 9 |
| 18 h | -5.388 | <0.01 | 9 |
| 24 h | 0.332 | 1 | 5 |
| **Fig. 2A, Standard protocol, paired t-test, Bonferroni correction** |  |  |  |
|  | t | p | n |
| 0 h | -4.362 | 0.042 | 6 |
| 1 h | -3.600 | 0.042 | 9 |
| 2 h | 0.851 | 1 | 7 |
| 5 h | -4.048 | 0.03 | 8 |
| 18 h | -5.483 | 0.018 | 6 |
| 24 h | -1.312 | 1 | 8 |
| **Fig. 2A, Enhanced protocol, paired t-test, Bonferroni correction** |  |  |  |
|  | t | p | n |
| 0 h | -5.138 | 0.042 | 5 |
| 1 h | -3.924 | 0.048 | 7 |
| 2 h | -0.544 | 1 | 5 |
| 5 h | -4.896 | 0.048 | 5 |
| 18 h | -5.342 | 0.006 | 8 |
| 24 h | -1.237 | 1 | 8 |
| **Fig. 2A, two-pulse protocol, paired t-test, Bonferroni correction** |  |  |  |
|  | t | p | n |
| 0 h | -4.741 | 0.03 | 6 |
| 1 h | -4.330 | 0.03 | 7 |
| 2 h | -0.0420 | 1 | 5 |
| 5 h | -3.919 | 0.048 | 7 |
| 18 h | -4.010 | 0.024 | 9 |
| 24 h | -0.447 | 1 | 8 |
| **Fig. 3A, two-pulse protocol, Wilcoxon Signed Rank Test, Bonferroni correction** |  |  |  |
|  | z | p | n |
| 0 h | -0.169 | 1 | 7 |
| 1 h | 2.521 | 0.048 | 8 |
| 2 h | -0.507 | 1 | 7 |
| 5 h | 2.666 | 0.024 | 9 |
| 18 h | 2.666 | 0.024 | 9 |
| 24 h | 0.178 | 1 | 9 |
| **Fig. 3A, Enhanced protocol, paired t-test, Bonferroni correction** |  |  |  |
|  | t | p | n |
| 0 h | 0.761 | 1 | 8 |
| 1 h | 3.564 | 9 | 9 |
| 2 h | 1.401 | 1 | 7 |
| 5 h | 3.714 | 0.036 | 9 |
| 18 h | 3.502 | 0.048 | 9 |
| 24 h | 3.919 | 0.036 | 8 |
| **Fig. 4B2 one-way RM ANOVA ($F_{3,33} = 5.073$, $p = 0.005$), pairwise comparisons (Student-Newman-Keuls)** |  |  |  |
|  | q | P | n |
| 5-HT alone vs. Veh group | 1.95 | 0.363 | 12 |
| 5-HT alone vs. BID alone group | 1.544 | 0.283 | 12 |
| 5-HT alone vs. 5-HT + BID group | 3.015 | 0.041 | 12 |
| BID alone vs. Veh group | 0.406 | 0.776 | 12 |
| 5-HT + BID vs. Veh group | 4.965 | 0.007 | 12 |
| BID alone vs. 5-HT + BID group | 4.559 | 0.008 | 12 |
| **Fig. 4C2 Friedman repeated measures analysis of variance on ranks (chi-square = 11.850, with 3 degrees of freedom, $p = 0.008$), pairwise comparisons (Student-Newman-Keuls)** |  |  |  |
|  | q | P | n |
| 5-HT alone vs. Veh group | 2.500 | >0.05 | 8 |
| 5-HT alone vs. SB alone group | 3.536 | <0.05 | 8 |
| 5-HT alone vs. 5-HT + SB group | 3.500 | <0.05 | 8 |
| SB alone vs. Veh group | 2.500 | >0.05 | 8 |
| 5-HT + SB vs. Veh group | 4.243 | <0.05 | 8 |
| SB alone vs. 5-HT + SB group | 4.656 | <0.05 | 8 |
| **Fig. 5B2 one-way RM ANOVA revealed a significant overall effect of the treatments ($F_{3,18} = 6.338$, $p = 0.004$), pairwise comparisons (Student-Newman-Keuls)** |  |  |  |
|  | q | P | n |
| 5-HT alone vs. Veh group | 5.98 | 0.003 | 7 |
| 5-HT alone vs. KT5720 alone group | 3.971 | 0.012 | 7 |
| 5-HT alone vs. 5-HT + KT5720 group | 4.127 | 0.024 | 7 |
| KT5720 alone vs. Veh group | 2.009 | 0.352 | 7 |
| 5-HT + KT5720 vs. Veh group | 1.853 | 0.207 | 7 |
| KT5720 alone vs. 5-HT + KT5720 group | 0.156 | 0.914 | 7 |
| **Fig. 5C2 one-way RM ANOVA revealed a significant overall effect of the treatments ($F_{3,12} = 7.61$, $p = 0.004$), pairwise comparisons (Student-Newman-Keuls)** |  |  |  |
|  | q | P | n |
| 5-HT alone vs. Veh group | 5.425 | 0.006 | 5 |
| 5-HT alone vs. KT5720 alone group | 5.343 | 0.003 | 5 |
| 5-HT alone vs. 5-HT + KT5720 group | 5.751 | 0.007 | 5 |
| KT5720 alone vs. Veh group | 0.082 | 0.955 | 5 |
| 5-HT + KT5720 vs. Veh group | 0.326 | 0.822 | 5 |
| KT5720 alone vs. 5-HT + KT5720 group | 0.408 | 0.955 | 5 |
| **Fig. 6B2 Friedman repeated measures analysis of variance on ranks (chi-square=10.029, with 3 degrees of freedom, $p = 0.018$), pairwise comparisons (Student-Newman-Keuls)** |  |  |  |
|  | q | P | n |
| 5-HT alone vs. Veh group | 4.099 | < 0.05 | 7 |
| 5-HT alone vs. U0126 alone group | 3.402 | < 0.05 | 7 |
| 5-HT alone vs. 5-HT + U0126 group | 1.604 | > 0.05 | 7 |
| U0126 alone vs. Veh group | 2.673 | > 0.05 | 7 |
| 5-HT + U0126 vs. Veh group | 4.158 | < 0.05 | 7 |
| U0126 alone vs. 5-HT + U0126 group | 3.207 | < 0.05 | 7 |
| **Fig. 6C2 one-way RM ANOVA revealed a significant overall effect of the treatments ($F_{3,21} = 5.165$, $p = 0.008$), pairwise comparisons (Student-Newman-Keuls)** |  |  |  |
|  | q | P | n |
| 5-HT alone vs. Veh group | 5.38 | 0.005 | 8 |
| 5-HT alone vs. KT5720 alone group | 3.9 | 0.03 | 8 |
| 5-HT alone vs. 5-HT + KT5720 group | 3.34 | 0.028 | 8 |
| KT5720 Fc alone vs. Veh group | 1.48 | 0.307 | 8 |
| 5-HT + KT5720 vs. Veh group | 2.04 | 0.338 | 8 |
| KT5720 alone vs. 5-HT + KT5720 group | 0.561 | 0.696 | 8 |
| **Fig. 6D2 one-way RM ANOVA revealed a significant overall effect of the treatments ($F_{3,24} = 3.728$, $p = 0.025$), pairwise comparisons (Student-Newman-Keuls)** |  |  |  |
|  | q | P | n |
| 5-HT alone vs. Veh group | 3.432 | 0.023 | 9 |
| 5-HT alone vs. TrkB Fc alone group | 3.655 | 0.042 | 9 |
| 5-HT alone vs. 5-HT + TrkB Fc group | 4.290 | 0.027 | 9 |
| TrkB Fc alone vs. Veh group | 0.223 | 0.876 | 9 |
| 5-HT + TrkB Fc vs. Veh group | 0.858 | 0.818 | 9 |
| TrkB Fc alone vs. 5-HT + TrkB Fc group | 0.635 | 0.658 | 9 |
| **Fig. 7B2 one-way RM ANOVA revealed a significant overall effect of the treatments ($F_{3,30} = 5.059$, $p = 0.006$), pairwise comparisons (Student-Newman-Keuls)** |  |  |  |
|  | q | P | n |
| 5-HT alone vs. Veh group | 4.973 | 0.007 | 11 |
| 5-HT alone vs. TGF-β RII Fc alone group | 4.499 | 0.009 | 11 |
| 5-HT alone vs. 5-HT + TGF-β RII Fc group | 3.543 | 0.018 | 11 |
| TGF-β RII Fc alone vs. Veh group | 0.474 | 0.74 | 11 |
| 5-HT + TGF-β RII Fc vs. Veh group | 1.43 | 0.576 | 11 |
| TGF-β RII Fc alone vs. 5-HT + TGF-β RII Fc group | 0.956 | 0.504 | 11 |
| **Fig. 7C2 one-way RM ANOVA revealed a significant overall effect of the treatments ($F_{3,36} = 5.037$, $p = 0.005$), pairwise comparisons (Student-Newman-Keuls)** |  |  |  |
|  | q | P | n |
| 5-HT alone vs. Veh group | 4.569 | 0.007 | 13 |
| 5-HT alone vs. TGF-β RII Fc alone group | 4.914 | 0.007 | 13 |
| 5-HT alone vs. 5-HT + TGF-β RII Fc group | 2.87 | 0.05 | 13 |
| TGF-β RII Fc alone vs. Veh group | 0.345 | 0.809 | 13 |
| 5-HT + TGF-β RII Fc vs. Veh group | 1.699 | 0.238 | 13 |
| TGF-β RII Fc alone vs. 5-HT + TGF-β RII Fc group | 2.043 | 0.329 | 13 |
| **Fig. 7D2 one-way RM ANOVA revealed a significant overall effect of the treatments ($F_{3,23} = 4.39$, $p = 0.014$), pairwise comparisons (Student-Newman-Keuls)** |  |  |  |
|  | q | P | n |
| 5-HT alone vs. Veh group | 4.234 | 0.017 | 8 |
| 5-HT alone vs. TGF-β RII Fc alone group | 4.492 | 0.021 | 8 |
| 5-HT alone vs. 5-HT + TGF-β RII Fc group | 4.027 | 0.009 | 8 |
| TGF-β RII Fc alone vs. Veh group | 0.268 | 0.851 | 9 |
| 5-HT + TGF-β RII Fc vs. Veh group | 0.216 | 0.88 | 9 |
| TGF-β RII Fc alone vs. 5-HT + TGF-β RII Fc group | 0.484 | 0.938 | 9 |
| **Fig. 8B2 one-way RM ANOVA revealed a significant overall effect of the treatments ($F_{3,15} = 4.675$, $p = 0.017$), pairwise comparisons (Student-Newman-Keuls)** |  |  |  |
|  | q | P | n |
| 5-HT alone vs. Veh group | 3.856 | 0.039 | 6 |
| 5-HT alone vs. TGF-β RII Fc alone group | 3.284 | 0.035 | 6 |
| 5-HT alone vs. 5-HT + TGF-β RII Fc group | 5.055 | 0.013 | 6 |
| TGF-β RII Fc alone vs. Veh group | 0.571 | 0.692 | 6 |
| 5-HT + TGF-β RII Fc vs. Veh group | 1.199 | 0.410 | 6 |
| TGF-β RII Fc alone vs. 5-HT + TGF-β RII Fc group | 1.770 | 0.443 | 6 |
| **Fig. 8C2 one-way RM ANOVA revealed a significant overall effect of the treatments ($F_{3,16} = 5.189$, $p = 0.011$), pairwise comparisons (Student-Newman-Keuls)** |  |  |  |
|  | q | P | n |
| 5-HT alone vs. Veh group | 5.244 | 0.009 | 7 |
| 5-HT alone vs. SIS3 alone group | 3.442 | 0.027 | 6 |
| 5-HT alone vs. 5-HT + SIS3 group | 4.08 | 0.028 | 6 |
| SIS3 alone vs. Veh group | 1.532 | 0.538 | 6 |
| 5-HT + SIS3 vs. Veh group | 0.894 | 0.536 | 6 |
| SIS3 alone vs. 5-HT + SIS3 group | 0.608 | 0.673 | 6 |
| **Fig. 8D2 one-way RM ANOVA revealed a significant overall effect of the treatments ($F_{3,21} = 5.337$, $p = 0.007$), pairwise comparisons (Student-Newman-Keuls)** |  |  |  |
|  | q | P | n |
| 5-HT alone vs. Veh group | 5.124 | 0.008 | 8 |
| 5-HT alone vs. SIS3 alone group | 4.641 | 0.01 | 8 |
| 5-HT alone vs. 5-HT + SIS3 group | 3.277 | 0.031 | 8 |
| SIS3 alone vs. Veh group | 0.483 | 0.736 | 8 |
| 5-HT + SIS3 vs. Veh group | 1.846 | 0.408 | 8 |
| SIS3 alone vs. 5-HT + SIS3 group | 1.364 | 0.346 | 8 |

At the end of the 60-min Enhanced protocol, p-p38 MAPK was increased to 37.9 ± 13.7% of control. At 1 h it was increased to 34.5 ± 6.3% of control, but declined 2 h (3.5 ± 7.0%). p-p38 MAPK increased again at 5 h (23.1 ± 8.0%) and 18 h (25.1 ± 7.5%), followed by a decay at 24 h (17.6 ± 10.3%). Following the 50-min two-pulse protocol, p-p38 MAPK increased at 0 h (41.8 ± 10.0%), and 1 h (21.3 ± 9.7%), but declined at 2 h (4.1 ± 12.8%). The second wave of increase of p-p38 MAPK was evident at 5 h (36.7 ± 15.8%), and 18 h (44.8 ± 10.2%), followed by a return to basal level at 24 h (−3.1 ± 12.3%).

Because all three protocols induced similar dynamics at early times, we focused on the comparison of p-p38 MAPK at 18 h and 24 h induced by the three protocols (Fig. 1B-C), using the same data as in Fig. 1A. Although p-p38 MAPK at 24 h after the Enhanced protocol appeared to remain elevated, statistical analyses (one way ANOVA) did not reveal significant differences between p-p38 MAPK at 18 h (Fig. 1B) or 24 h (Fig. 1C) induced by the protocols (18 h, F_2,25_ = 1.479, p = 0.247; 24 h, F_2,20_ = 1.192, p = 0.324). Thus, despite differences in the duration of the protocols, and in effectiveness in inducing LTF (Zhang et al. 2011, 2023), the three protocols produced remarkably similar changes in the dynamics of p-p38 MAPK.

#### 2. Two waves of increase in PKA activity (PKA catalytic subunits) and in RSK activity (pRSK) are also induced within 24h

PKA phosphorylates CREB1 (Bartsch et al. 1998) and also contributes to the activity of RSK that, in parallel with PKA, activates CREB1 (Zhang et al. 2021; Liu et al. 2022). To investigate the dynamics of PKA, the levels of catalytic subunits of PKA (PKAc), which is an active form of PKA, were measured at 0 h, 1 h, 2 h, 5 h, 18 h, and 24 h. The Standard (Fig. 2A, grey line), Enhanced (Fig. 2A, red line) and two-pulse (Fig. 2A, blue line) protocols all induced two waves of increase. After the Standard protocol, PKAc increased at 0 h (32.7 ± 10.8%), and 1 h (24.7 ± 6.7%), but declined at 2 h (−2.5 ± 10.0%). PKAc increased again at 5 h (16.3 ± 3.4%), the increase was maintained at 18 h (35.2 ± 10.7%), followed by a decrease toward basal level at 24 h (22.7 ± 12.5%), a value not statistically different from time-matched controls.

**Figure 2.**
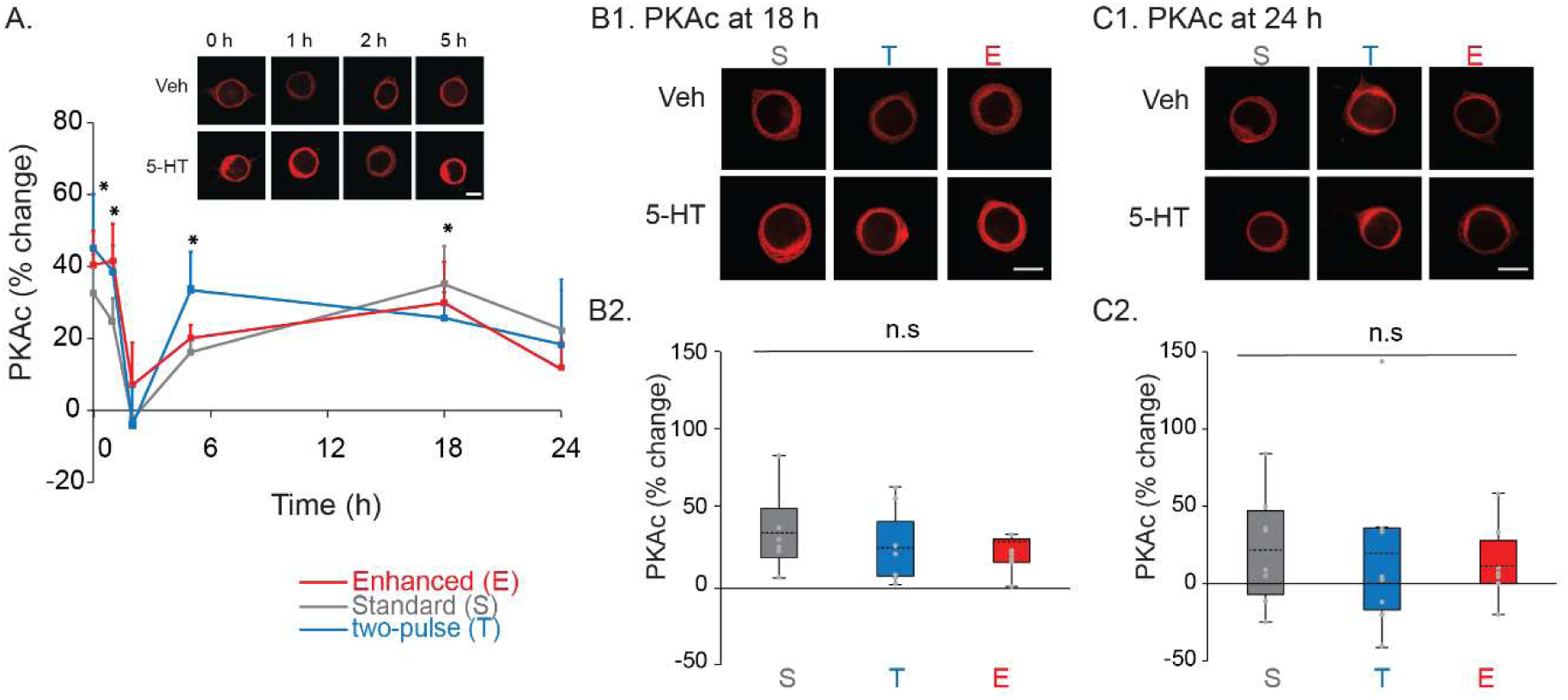
Dynamics of PKA catalytic subunits (PKAc) induced by three LTF-inducing protocols. ***A***, Time course of PKAc, with representative confocal images (top) and summary data (bottom). The percent change was calculated as the change of PKAc level after 5-HT compared to time-matched vehicle control. For all protocols, PKAc was increased immediately (0 h) after 5-HT and at 1 h after, returned to near basal levels at 2 h, and then increased again at 5 h. This increase was maintained at 18 h for all protocols, followed by a return toward basal level at 24 h. * p < 0.05. ***B***, Comparison of PKAc at 18 h after different 5-HT protocols. ***B1***, Representative confocal images of PKAc immunostaining in SNs. ***B2***, Summary data. Statistical analyses did not reveal significant differences between PKAc at 18 h induced by three protocols. ***C***, Comparison of PKAc at 24 h after different 5-HT protocols. ***C1***, Representative confocal images of PKAc immunostaining in SNs. ***C2***, Summary data. Statistical analyses did not reveal significant differences between PKAc at 24 h induced by three protocols.

The Enhanced protocol increased PKAc at 0 h (40.5 ± 9.6%), and 1 h (41.5 ± 9.9%), followed by a decline at 2 h (7.2 ± 11.3%) after treatment. PKAc increased again at 5 h (20.2 ± 3.6%), the increase was maintained at 18 h (29.9 ± 11.4%), followed by a decrease toward basal level at 24 h (11.5 ± 8.4%). After the two-pulse protocol, PKAc increased at 0 h (45.3 ± 15.1%), and 1 h (40.0 ± 7.9%), but declined at 2 h (−4.5 ± 10.8%). PKAc increased again at 5 h (33.5 ± 10.7%), the increase was maintained at 18 h (25.8 ± 7.2%), followed by a decrease toward basal level at 24 h (18.4 ± 20.1%).

PKAc levels at 18 h and 24 h were compared between the three protocols (Fig. 2B-C). Statistical analyses (Kruskal-Wallis one way analysis of variance on ranks) did not reveal significant differences between PKAc at 18 h (Fig. 2B) or 24 h (Fig. 2C) (18 h, H_2_ = 1.266, p = 0.531; 24 h, H_2_ = 0.980, p = 0.613). Thus, all three protocols induced similar long-term dynamics of PKAc.

Liu et al. (2019) identified a complex dynamic of phosphorylated RSK (pRSK) mediated by the Standard protocol, but did not examine the 18 h time point, nor examine the dynamics induced by the Enhanced and two-pulse protocols. We found that the Enhanced (Fig. 3A, red line) and two-pulse (Fig. 3A, blue line) protocols induced two waves of increase in pRSK. After the two-pulse protocol, pRSK increased at 1 h (22.2 ± 6.4%), but not at 0 h (−0.2 ± 8.6%), or 2 h (−3.5 ± 10.4%). pRSK increased again at 5 h (40.4 ± 12.0%), the increase was maintained at 18 h (36.4 ± 13.1%), followed by a return to basal level at 24 h (2.2 ± 9.2%). After the Enhanced protocol, pRSK increased at 1 h (20.5 ± 6.0%), but not at 0 h (−0.6 ± 15.9%), or 2 h (−7.5 ± 7.6%). pRSK increased again at 5 h (23.1 ± 5.9%), the increase was maintained at 18 h (30.2 ± 8.6%), and, in contrast to the Standard and two-pulse groups, at 24 h (44.9 ± 12.9%). We also measured pRSK at 18 h after the Standard protocol, a time point not assessed previously (Fig. 3B). pRSK significantly increased by 23.1 ± 3.6% (n = 8, Wilcoxon Signed Rank Test, z = 2.521, p = 0.048). Statistical analyses did not reveal any significant difference between pRSK at 18 h induced by three protocols (Fig. 3B2) (Kruskal-Wallis one way analysis of variance on ranks, H_2_ = 0.308, p = 0.857).

**Figure 3.**
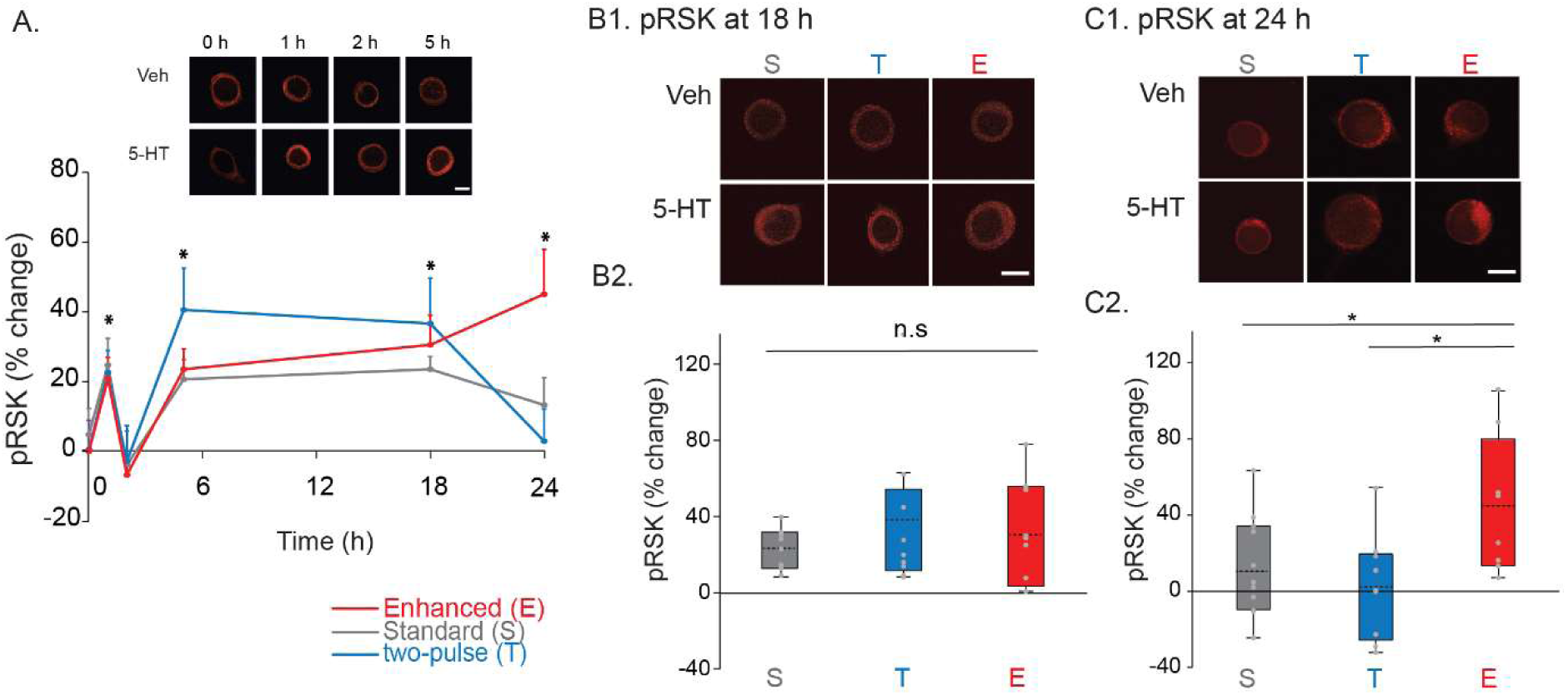
Dynamics of pRSK induced by three LTF-inducing protocols. ***A***, Time course of RSK activity, with representative confocal images (top) and summary data (bottom). The percent change was calculated as the change of pRSK level after 5-HT compared to time-matched vehicle control. For all protocols, RSK was increased at 1 h after 5-HT, returned to near basal levels at 2 h, and then increased again at 5 h. This increase was maintained at 18 h for all protocols (data of Standard protocol are from Liu et al. 2019 except for 18 h). For the Standard and two-pulse protocols, RSK returned to basal at 24 h. However, pRSK induced by the Enhanced protocol remained elevated at 24 h (red). ***B***, Comparison of pRSK at 18 h after different 5-HT protocols. ***B1***, Representative confocal images of pRSK immunostaining in SNs. ***B2***, Summary data. Statistical analyses did not reveal significant differences between pRSK at 18 h induced by three protocols. ***C***, Comparison of pRSK at 24 h after different 5-HT protocols. ***C1***, Representative confocal images of pRSK immunostaining in SNs. ***C2***, Summary data. pRSK induced by the Enhanced protocol was significantly greater than pRSK induced by the other two protocols. * p < 0.05.

However, at 24 h (Fig. 3C), a one-way ANOVA revealed a significant overall effect of the treatments (F_2, 25_ = 4.608, p = 0.02). Subsequent pairwise comparisons (Student-Newman-Keuls) revealed that the Enhanced group was significantly different from the two-pulse group (q = 4.131, p = 0.019), and the Standard group (q = 3.261, p = 0.03). No significant difference was detected between the two-pulse group and the Standard group (q = 1.095, p = 0.446). These results suggest that the Enhanced protocol produces a unique dynamic signature in pRSK at 24 h.

The profiles of pRSK elicited by the three protocols are also distinct in that pRSK was initially undetectable immediately after the end of treatment and only increased 1 h later (Fig. 3A), whereas p-p38 MAPK (Fig. 1), PKAc (Fig. 2), and pERK (Zhang et al. 2023a) were elevated immediately after the end of treatment.

### II. Determinants of the Dynamic Signatures of the Kinases

We next sought to obtain insights into the processes that regulate the dynamic signatures of the kinases, in particular RSK. We tested the hypothesis that the profiles were due to a combination of feedback loops among the cascades and partially extracellular feedback loops. Previous studies have characterized feedback loops, with extracellular components, involving the assessed kinases and the growth factors *Aplysia* neurotrophin (NT) (Jin et al. 2018) and transforming growth factor-β (TGF-β) (Kopec et al. 2015).

#### 1. An inhibitory feedback loop was responsible for the delay in the first wave of increase of RSK activity after 5-HT treatment

A significant difference was observed between the early activation of p38 MAPK (Fig. 1), PKA (Fig. 2) and pERK (Zhang et al. 2023a), and the delayed activation of RSK (Fig. 3A) following the end of treatment. The delay in the activation of RSK was surprising because RSK is activated immediately after a single 5-min pulse of 5-HT (Zhang et al. 2021), and RSK is downstream of ERK (Liu et al. 2019), which is activated both during the application of 5-HT (Zhang et al. 2021) and immediately after the end of training (Zhang et al. 2023a), as are p-p38 MAPK (Fig. 1) and PKA (Fig. 2). Previous studies identified a negative feedback pathway from p38 MAPK to ERK after one pulse of 5-HT treatment (Fig. 4A, solid curves) (Zhang et al. 2017, 2021), however, this pathway by itself seems unlikely to delay RSK activation after 5 pulses of 5-HT, because pERK was significantly increased immediately at the end of treatment (Zhang et al. 2023a). Therefore, we considered the possibility that the delay in pRSK increase might be due to another negative feedback loop independent of ERK. One hypothesis is a pathway by which RSK activates p38 MAPK and p38 MAPK feeds back to inhibit RSK (Fig. 4A, dashed curve).

**Figure 4.**
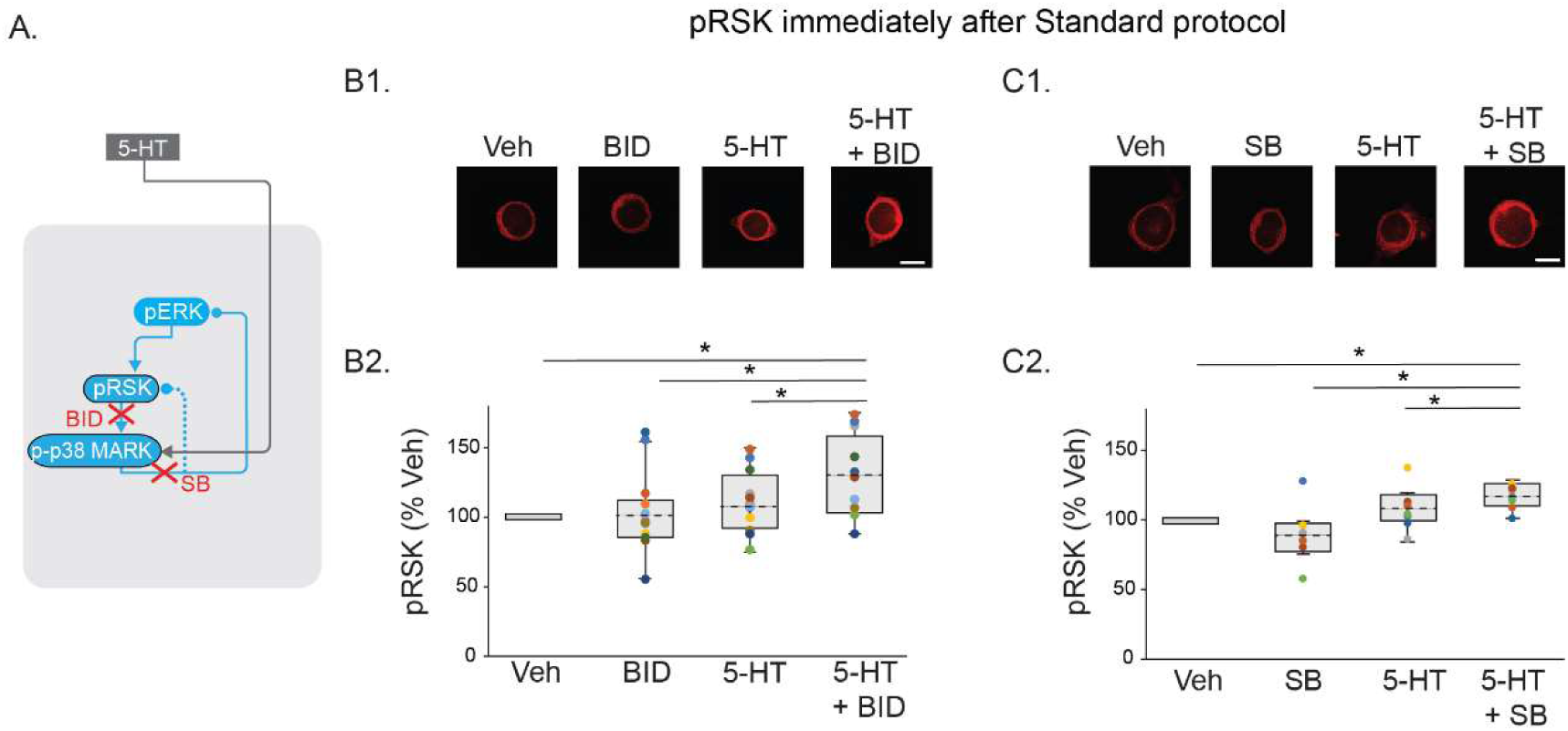
The repression in pRSK increase immediately after the Standard protocol was relieved by inhibition of RSK (***B***) or p38 MAPK (***C***). ***A***, Partial circuit diagram showing pathways targeted with inhibitors. ***B1***, Representative confocal images immediately after 5-HT, in the absence or presence of RSK inhibitor BI-D1870 (BID). ***B2***, Summary data. BID significantly increased pRSK induced by 5-HT. ***C1***, Representative confocal images of pRSK in SNs immediately after 5-HT, in the absence or presence of p38 MAPK inhibitor SB203580 (SB). ***C2***, Summary data. SB significantly increased pRSK induced by 5-HT. * p < 0.05. In ***B2*** and subsequent plots, circles with the same color indicate the results are from the same animal.

To test this hypothesis, we applied the RSK inhibitor BI-D1870 (BID), or the p38 MAPK inhibitor SB203580 (SB) for 0.5 h before and during the Standard protocol, and measured the level of pRSK immediately after 5-HT (Fig. 4B-C). For these and subsequent inhibitor experiments, four dishes of SNs from the same animal were used for each experiment: 1) 50 μM 5-HT alone; 2) inhibitor alone; 3) 5-HT + inhibitor; or 4) Veh alone (as described in Methods). Example responses are illustrated in Figs. 4B1, 4C1 and summary data in Figs. 4B2, 4C2. In the presence of BID (Fig. 4B), which inhibits the activity, but not the phosphorylation, of RSK, levels of pRSK increased (28.1 ± 8.4%), whereas, and replicating the results of Fig. 3, 5-HT did not induce a significant increase (3.9 ± 8.6%) in levels of pRSK immediately after treatment, in the absence of BID. A one-way RM ANOVA revealed a significant overall effect of the treatments (Table 1). Subsequent pairwise comparisons revealed that the 5-HT + BID group was significantly different from the other three groups. No significant differences were observed between the other groups. Therefore, RSK activity was significantly increased immediately after treatment if BID was applied before and during treatment, indicating RSK activates an auto-regulating feedback inhibition.

To test the involvement of p38 MAPK, the experiment was repeated in the presence of SB (Fig. 4C). 5-HT induced a significant increase (16.7 ± 2.8%) in pRSK, compared to the level in the absence of SB (7.8 ± 4.9%). Friedman repeated measures analysis of variance on ranks revealed a significant overall effect of the treatments (Table 1). Subsequent pairwise comparisons revealed that the 5-HT + SB group was significantly different from the other three groups. No significant differences were observed between the other groups. Thus, the mechanism underlying the delayed increase of pRSK immediately after 5-HT protocols might be as follows. pRSK is initially increased during the 5 pulses of 5-HT (Zhang et al. 2021). This increase then activates p38 MAPK, which then feeds back to reduce pRSK towards basal by the end of 5-HT treatment (Fig. 4A).

#### 2. The 5-HT-induced first wave of pERK and pRSK is PKA dependent

The first wave of PKA activity is due, at least in part, to an extracellular feedback loop involving the PKA-induced release of *Aplysia* neurotrophin (NT), and subsequent activation by NT of Trk receptors on the sensory neuron, which in turn further activates PKA and ERK (Jin et al. 2018; Zhang et al. 2021) (circuit diagram, Fig. 5A). To test whether this feedback loop also contributes to the first wave of increase of pERK and pRSK, we applied the PKA inhibitor KT5720 for 0.5 h before and during the treatment with the Standard protocol, and measured the level of pERK (Fig. 5B) and pRSK (Fig. 5C) at 1 h after 5-HT. Example responses are illustrated in Figs. 5B1, 5C1 and summary data are presented in Figs. 5B2, 5C2. 5-HT induced a significant increase (36.8 ± 10.2%) in levels of pERK at 1 h after treatment, but levels of pERK were reduced by KT5720 (14.3 ± 9.9%) (Fig. 5B). A one-way RM ANOVA revealed a significant overall effect of the treatments (Table 1). Subsequent pairwise comparisons revealed that the 5-HT alone group was significantly different from the other three groups. No significant differences were observed between the other groups.

**Figure 5.**
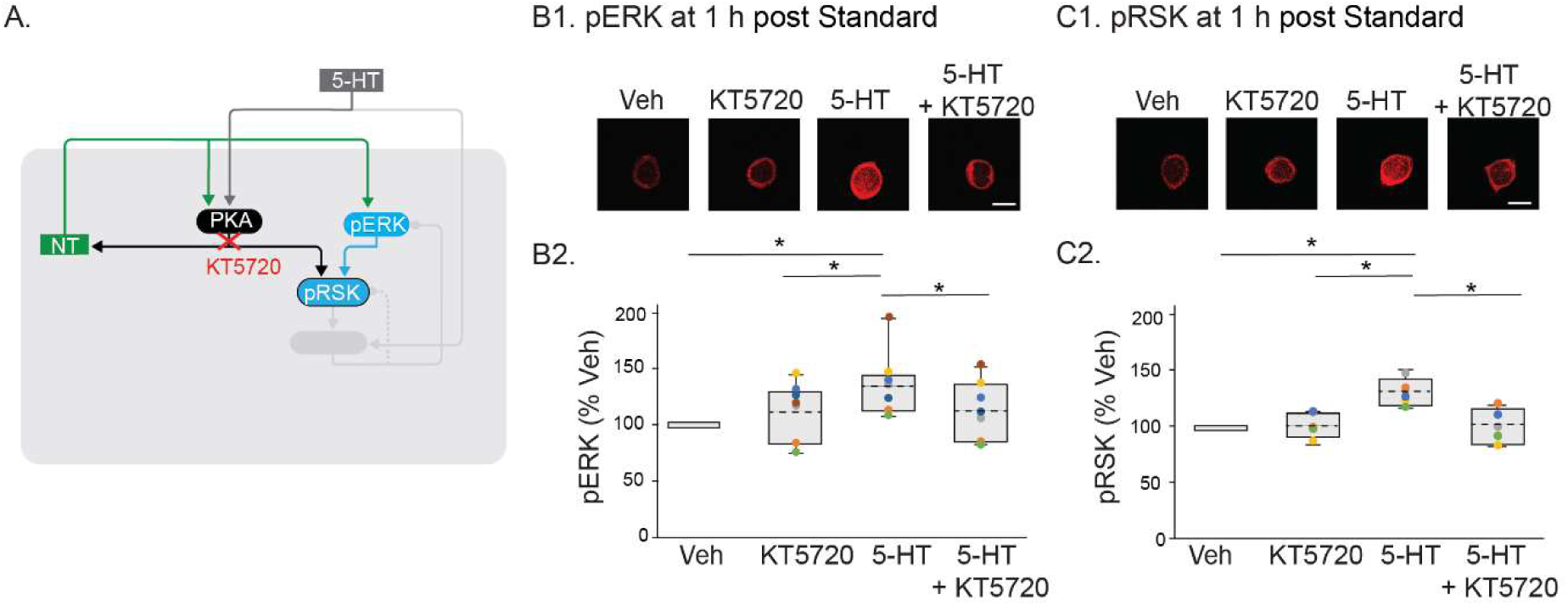
The first wave of increase in pRSK and pERK was dependent on PKA activity during the induction phase of LTF. ***A,*** Partial circuit diagram showing pathways targeted with inhibitors. ***B1***, Representative confocal images of pERK in SNs at 1 h after Standard protocol of 5-HT, in the absence or presence of PKA inhibitor KT5720. ***B2***, Summary data. KT5720 significantly decreased pERK induced by 5-HT. ***C1***, Representative confocal images of pRSK in SNs at 1 h after 5-HT, in the absence or presence of PKA inhibitor KT5720. ***C2***, Summary data. KT5720 significantly decreased pRSK induced by 5-HT. * p < 0.05.

Similarly, whereas 5-HT induced a significant increase (29.2 ± 5.1%) in levels of pRSK at 1 h after treatment (Fig. 5C), pRSK was reduced by KT5720 (0.5 ± 6.6%). A one-way RM ANOVA revealed a significant overall effect of the treatments (Table 1). Subsequent pairwise comparisons revealed that the 5-HT alone group was significantly different from the other three groups. No significant differences were observed between the other groups. Therefore, the activation of the ERK-RSK pathway during the first wave of increase was dependent on PKA activity (Fig. 5A).

#### 3. The initiation of the second wave of increase of pRSK was regulated by multiple pathways, likely independent of ERK

The level of pRSK significantly increased at 5 h after the Standard protocol (Fig. 3A). This could be due to an upstream upregulation of pERK. However, previous study did not detect a significant increase in pERK at this time point (Zhang et al. 2023a). Thus, it appears that the second wave of increase of pRSK was not initiated by ERK. But we cannot exclude the possibility that some low level of pERK could be capable of mediating RSK activation. To test this hypothesis, we measured pRSK at 5 h with or without an inhibitor of ERK activation, U0126 (Fig. 6A). 5-HT led to an increase (61.6 ± 21.3%) in levels of pRSK (Fig. 6B). This increase was not significantly reduced (53.9 ± 29.4%) by U1026. Friedman repeated measures analysis of variance on ranks revealed a significant overall effect of the treatments (Table 1). Subsequent pairwise comparisons revealed that the 5-HT alone or 5-HT + U0126 group was significantly different from the other two groups. No significant differences were observed between the 5-HT alone and 5-HT+U1026 groups. Thus, U1026 did not block the increase of pRSK at 5 h after the Standard protocol, indicating the initiation of the second wave of increase of pRSK is independent of the ERK pathway.

**Figure 6.**
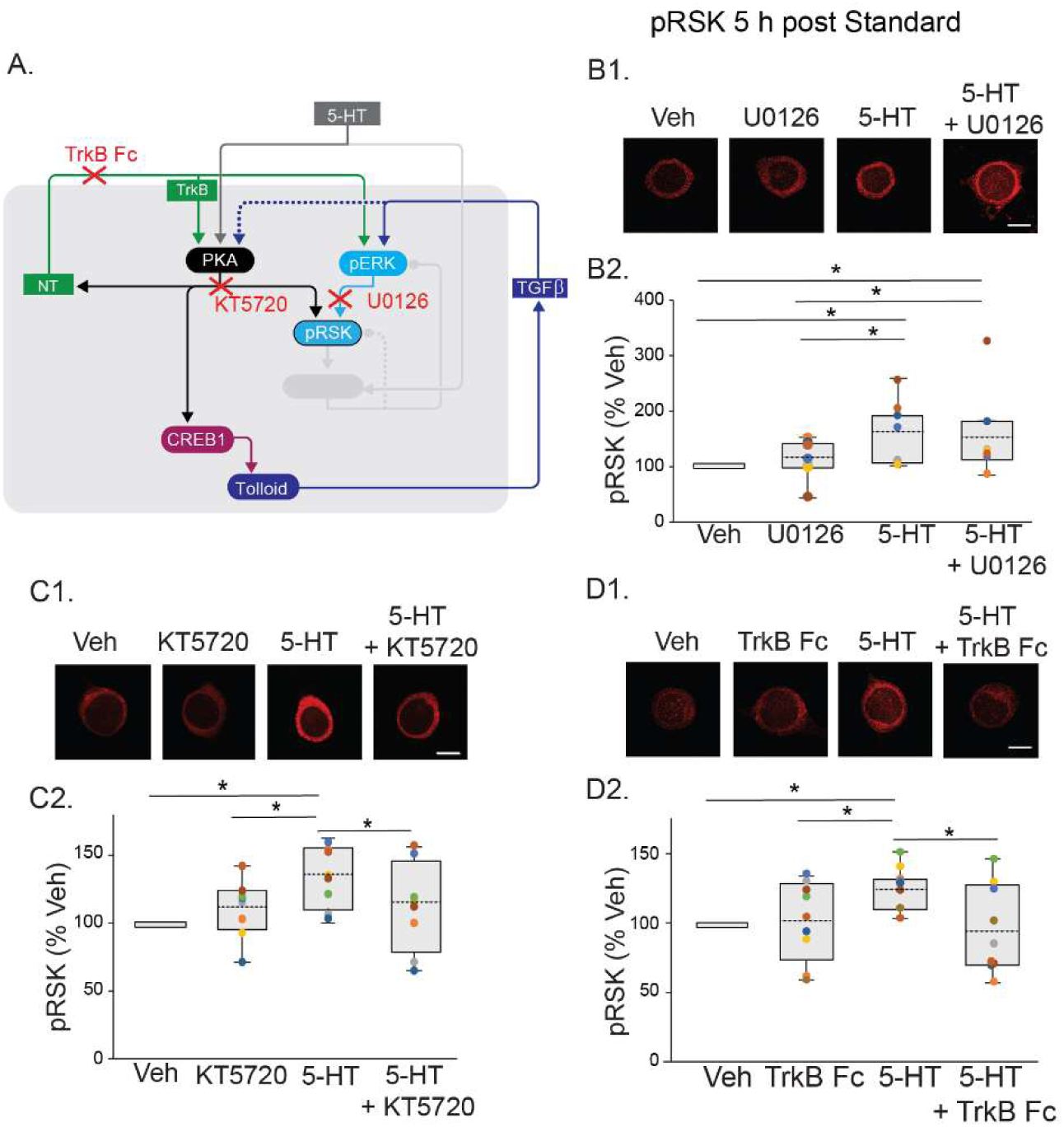
The second wave of increase in pRSK was repressed by inhibition of PKA or TrkB, but not by inhibition of ERK. ***A***, Partial circuit diagram showing pathways targeted with inhibitors. ***B1***, Representative confocal images of pRSK in SNs at 5 h after Standard protocol of 5-HT, in the absence or presence of inhibitor U0126. ***B2***, Summary data. U0126 did not reduce pRSK induced by 5-HT. ***C1***, Representative confocal images of pRSK in SNs at 5 h after 5-HT, in the absence or presence of PKA inhibitor KT5720. ***C2***, Summary data. KT5720 significantly decreased pRSK induced by 5-HT. ***D1***, Representative confocal images of pRSK in SNs at 5 h after Standard protocol of 5-HT, in the absence or presence of TrkB inhibitor TrkB Fc. ***D2***, Summary data. TrkB Fc significantly decreased pRSK induced by 5-HT. * p < 0.05.

Two other hypotheses are that the increase in pRSK at 5 h could be due to a PKA-NT/Trk positive feedback loop (Jin et al. 2018; Zhang et al. 2021) (Fig. 6A) or/and a TGF-β positive feedback loop (Zhang et al. 1997, 2023; Kopec et al. 2015) (Fig. 7A). For these feedback loops, pRSK is not a direct loop component, but may be increased by branching reactions (Figs. 6A, 7A). Three inhibitors were used to test these hypotheses: the PKA inhibitor KT5720 (Fig. 6C), the Trk antagonist TrkB Fc (Fig. 6D), and the TGF-β antagonist TGF-β RII Fc (Fig. 7). Each inhibitor was applied to SNs 4 h post-5-HT. The cells were fixed at 5 h for immunofluorescence analysis.

**Figure 7.**
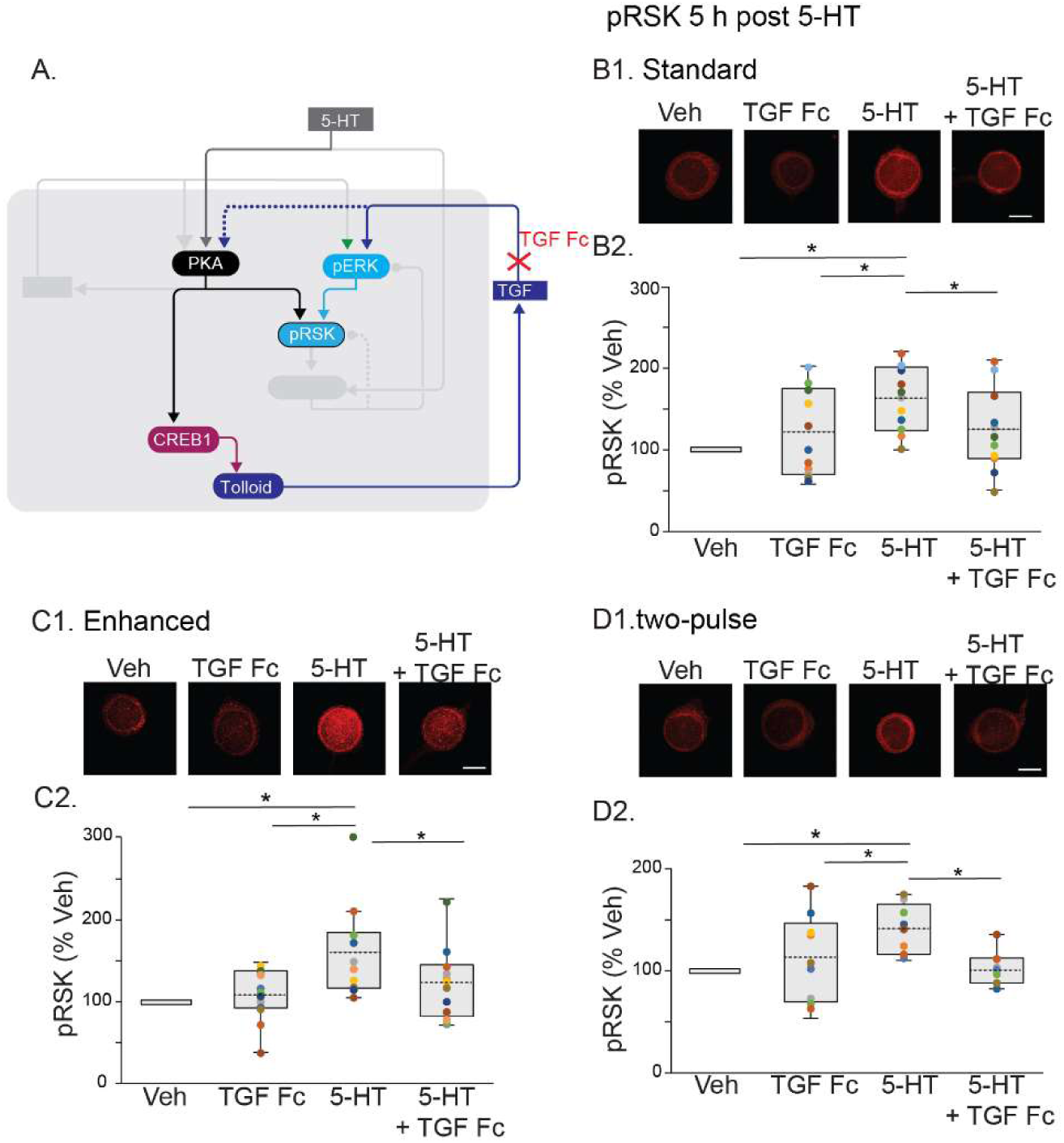
The second wave of increase in pRSK was repressed by inhibition of TGF-β. ***A***, Partial circuit diagram showing the pathway targeted by the TGF-β antagonist TGF-β RII Fc (TGF Fc). ***B1***, Representative confocal images of pRSK in SNs at 5 h after Standard protocol of 5-HT, in the absence or presence of TGF Fc. ***B2***, Summary data. TGF Fc significantly decreased pRSK induced by 5-HT. ***C1***, Representative confocal images of pRSK in SNs at 5 h after Enhanced protocol of 5-HT, in the absence or presence of TGF Fc. ***C2***, Summary data. TGF Fc significantly decreased pRSK induced by 5-HT. ***D1***, Representative confocal images of pRSK in SNs at 5 h after two-pulse protocol of 5-HT, in the absence or presence of TGF Fc. ***D2***, Summary data. TGF Fc significantly decreased pRSK induced by 5-HT. * p < 0.05.

##### PKA pathway

5-HT led to an increase (33.3 ± 7.6%) in levels of pRSK (Fig. 6C). This increase was blocked (11.6 ± 11.7%) by KT5720. A one-way RM ANOVA revealed a significant overall effect of the treatments (Table 1). Subsequent pairwise comparisons revealed that the 5-HT alone group was significantly different from the other three groups. No significant differences were observed between the other groups. Thus, the PKA pathway is involved in the second wave of increase in RSK activity (Fig. 6A).

##### Trk pathway

5-HT led to an increase (25.9 ± 5.2%) in levels of pRSK (Fig. 6D). This increase was blocked (−4.6 ± 10.5%) in the presence of TrkB Fc. A one-way RM ANOVA revealed a significant overall effect of the treatments (Table 1). Subsequent pairwise comparisons revealed that the 5-HT alone group was significantly different from the other three groups. No significant differences were observed between the other groups. Together with the results in Fig. 6C, these data further confirmed the of the second wave of increase in RSK activity was, at least partially, dependent on the PKA-NT/Trk pathway (Fig. 6A).

##### TGF-β pathway

5-HT led to an increase (51.1 ± 11.4%) in levels of pRSK at 5 h after the Standard protocol (Fig. 7B). This increase was suppressed (20.7 ± 13.8%) by the inhibitor of TGF-β, TGF-β RII Fc, which is chimeric protein sequestering TGF-β ligand. A one-way RM ANOVA revealed a significant overall effect of the treatments (Table 1). Subsequent pairwise comparisons revealed that the 5-HT alone group was significantly different from the other three groups. No significant differences were observed between the other groups. Thus, TGF-β pathway also contributed to the second wave of increase in pRSK after the Standard protocol.

We also examined the role of the TGF-β pathway in the activation of RSK by the Enhanced protocol, a strong LTF-inducing protocol. 5-HT led to an increase (53.2 ± 15.3%) in levels of pRSK (Fig. 7C). This increase was suppressed (19.7 ± 12.0%) by TGF-β RII Fc. A one-way RM ANOVA revealed a significant overall effect of the treatments (Table 1). Subsequent pairwise comparisons revealed that the 5-HT alone group was significantly different from the other three groups. No significant differences were observed between the other groups. Thus, the TGF-β pathway also contributed to the second wave of increase in RSK activity after the Enhanced protocol. Finally, the two-pulse protocol led to an increase (42.6 ± 8.5%) in levels of pRSK (Fig. 7D). This increase was blocked (1.9 ± 5.8%) by TGF-β RII Fc. A one-way RM ANOVA revealed a significant overall effect of the treatments (Table 1). Subsequent pairwise comparisons revealed that the 5-HT alone group was significantly different from the other three groups. No significant differences were observed between the other groups. Thus, the TGF-β pathway also contributed to the second wave of increase in RSK activity after this protocol (Fig. 7A).

As noted above, pRSK was elevated 5 h after the Standard protocol whereas pERK was not. Thus, although activation of RSK by ERK is established in other contexts, these data raised the interesting question of how RSK could be activated by TGF-β independent of ERK. One possibility is through a TGF-β-Smad-PKA pathway (Zhang et al. 2004; Miranda et al. 2023) (Fig. 8A).

**Figure 8.**
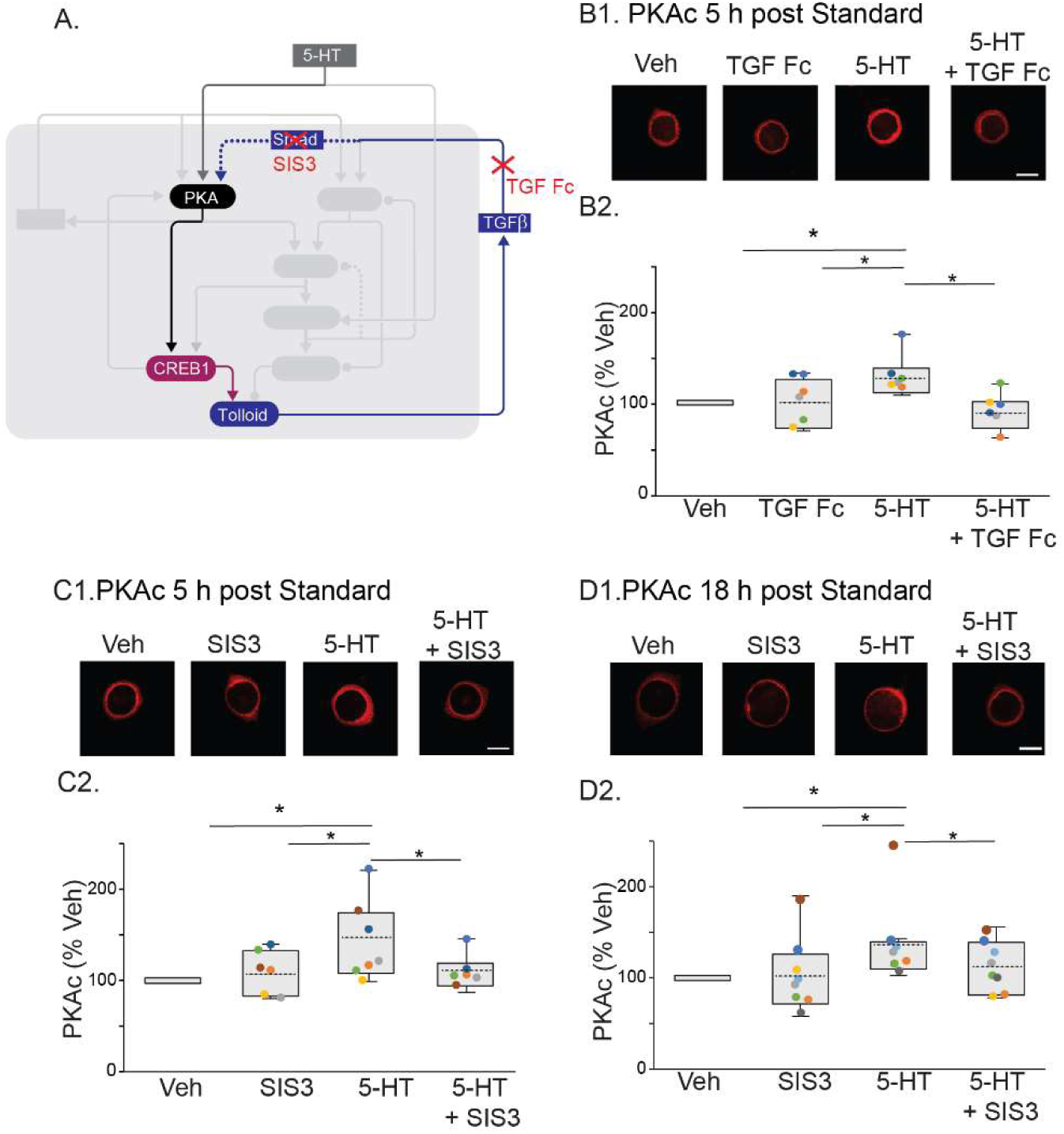
The second wave of increase in PKAc was repressed by inhibitors of TGF-β (**B**) or Smad3 (***C-D***). ***A***, Partial circuit diagram showing pathways targeted with inhibitors. ***B1***, Representative confocal images of PKAc in SNs 5 h after Standard protocol of 5-HT, in the absence or presence of TGF-β inhibitor TGF-β RII Fc (TGF Fc). ***B2***, Summary data. TGF Fc significantly decreased PKAc induced by 5-HT. ***C1***, Representative confocal images of PKAc in SNs 5 h after 5-HT, in the absence or presence of Smad3 inhibitor SIS3. ***C2***, Summary data. ***D1***, Representative confocal images of PKAc in SNs 18 h after 5-HT, in the absence or presence of Smad3 inhibitor SIS3. ***D2***, Summary data. SIS3 significantly decreased PKAc induced by 5-HT. * p < 0.05.

#### 4. The initiation of the second wave of increase of PKA activity was regulated by a TGF-**β**-SMAD3 pathway

The Smad protein family plays an important role in memory formation (Caraci et al. 2015; Gradari et al. 2021; Yu et al. 2014). TGFβ-R phosphorylates and activates the R-Smad protein in *Aplysia* (Miranda et al. 2023), the analog of Smad3 in mammalian cells (Zhang et al. 2004). In *Aplysia*, phosphorylated R-Smad forms a complex with co-Smad. This complex translocates to the nucleus and acts as a transcription cofactor (Miranda et al. 2023). In mammalian cells, phosphorylated Smad3 forms a complex with Smad4, which binds the regulatory subunits of PKA and releases free, active catalytic subunits (Zhang et al. 2004). We hypothesized that *Aplysia* R-Smad has an additional, similar function, directly activating PKA, forming a third feedback loop that contributes to the persistent increase of PKA. In this manner, the TGF-β pathway, activated by CREB1 (Zhang et al. 1997), may indirectly regulate the activity of PKA via R-Smad.

To test this hypothesis, we applied inhibitors of TGF-β and of Smad3 (Fig. 8A) to examine whether they suppressed the late increase of PKAc at 5 h and 18 h after the Standard protocol (Fig. 8B-D). The increase (32.0 ± 8.7%) in levels of PKAc at 5 h post-5HT (Fig. 8B) was indeed suppressed (−6.7 ± 7.9%) by TGF-β RII Fc. A one-way RM ANOVA revealed a significant overall effect of the treatments (Table 1). Subsequent pairwise comparisons revealed that the 5-HT alone group was significantly different from the other three groups. No significant differences were observed between the other groups. Therefore, the activation of the second wave of increase in PKAc after the Standard protocol was dependent on the TGF-β pathway.

The 5-HT-induced increase (43.6 ± 16.7%) in levels of PKAc at 5 h post 5-HT (Fig. 8C) was also suppressed (11.4 ± 7.2%) by the Smad3 inhibitor SIS3, thus SIS3 appears to inhibit R-Smad. A one-way RM ANOVA revealed a significant overall effect of the treatments (Table 1). Subsequent pairwise comparisons revealed that the 5-HT alone group was significantly different from the other three groups. No significant differences were observed between the other groups.

Similarly, at 18 h, the 5-HT-induced increase (38.4 ± 16.1%) in levels of PKAc at 18 h post 5-HT (Fig. 8D) was suppressed (12.8 ± 9.7%) by SIS3. A one-way RM ANOVA revealed a significant overall effect of the treatments (Table 1). Subsequent pairwise comparisons revealed that the 5-HT alone group was significantly different from the other three groups. No significant differences were observed between the other groups. Therefore, the second wave of increase in PKA activity after the Standard protocol was dependent on the PKA-CREB1-TGF-β/Smad pathway, with PKA in turn activating RSK (Fig. 8A).

## Discussion

### The differential dynamics of molecular pathways during 24 h after 5-HT protocols

Studies of molecular mechanisms underlying long-term synaptic plasticity typically focus on the changes of molecular pathways at a few specific times (e.g., immediately or 1 day after training). In this study, we systematically examined and compared the temporal changes of activities of essential kinases involved in consolidating LTF after three 5-HT protocols (Fig. 9).

**Figure 9.**
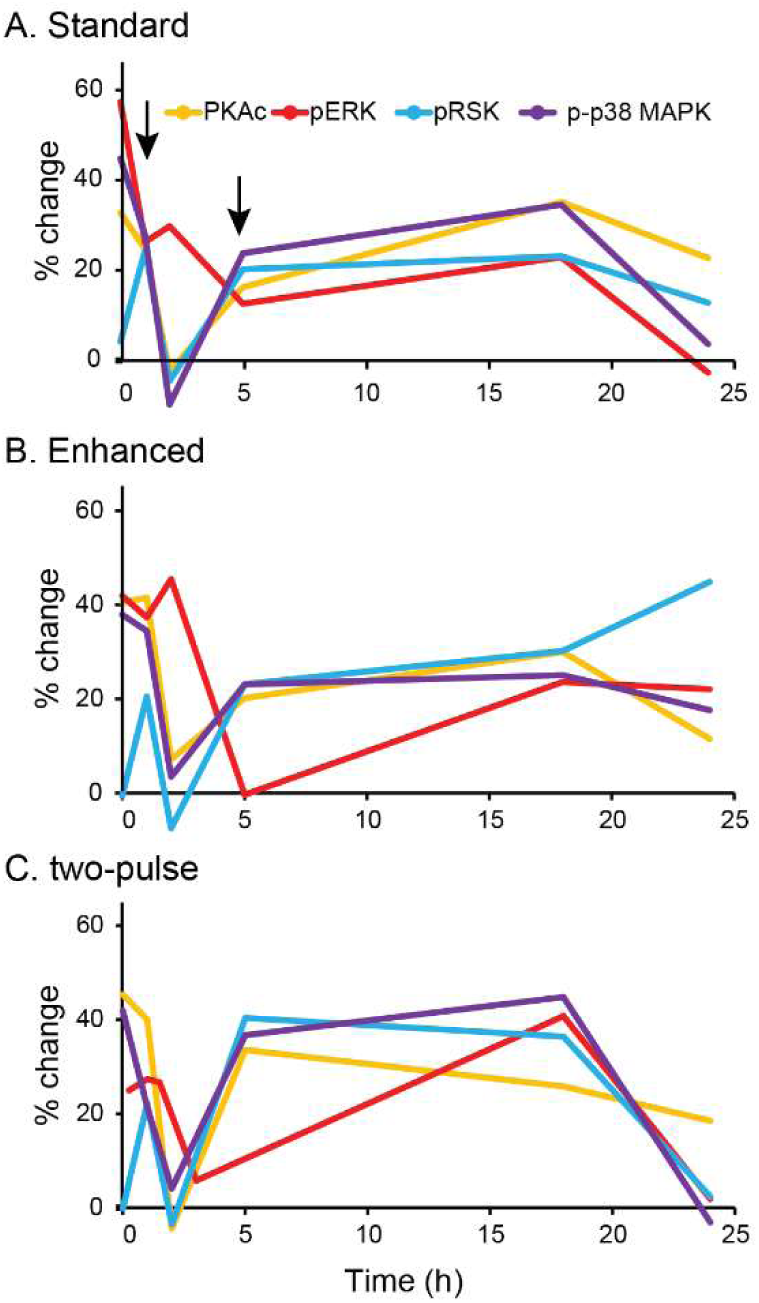
Dynamics of kinase activities after three protocols, combining results of Figs. 1-3, Liu et al. (2019) and Zhang et al. (2023). (For the two-pulse protocol, pERK was examined at 3 h and not 5 h.)

These protocols have substantial differences in the total amount of 5-HT application and duration of treatment. Despite these differences, the dynamics of activity of PKA, ERK, p38 MAPK, and RSK all showed two waves of increase after all three protocols. PKAc, p-p38 MAPK and pRSK had the first wave within 2 h of 5-HT application and the second lasting from ∼ 5 h to 18 h (arrows, Fig. 9). pERK had the first wave within 5 h and second wave near 18 h. But there were some key differences. pERK and pRSK returned towards basal at 24 h with the Standard and two-pulse protocols, but remained elevated at 24 h with the Enhanced protocol. There were also differences in the initial increase of activity. pRSK showed a delayed increase in all three protocols, whereas p-p38 MAPK, PKAc and pERK were elevated immediately after treatment.

### Complex interactions among molecular pathways activated by 5-HT

Figure 10 proposes a simplified model scheme of the signaling pathways activated by 5-HT. The results of Figs. 4-8 suggest that these interactions and interlocked feedback among kinases, transcription factors and growth factors, are critical for the waves of increases in kinase activity.

**Figure 10.**
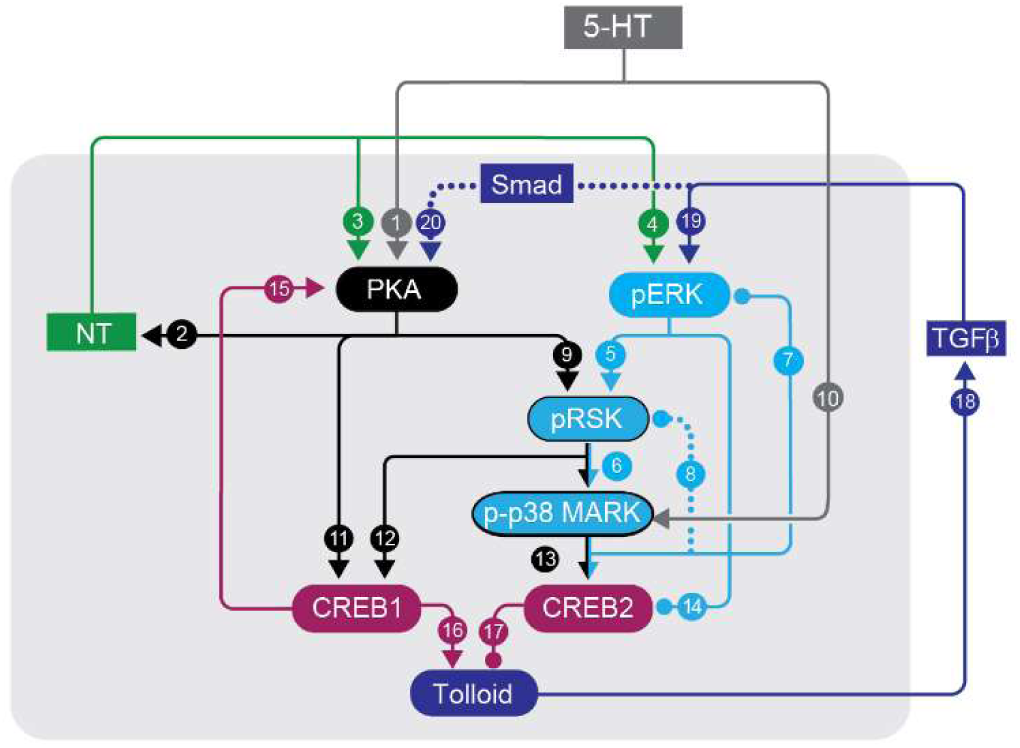
Schematic model of kinase signaling pathways critical for induction and consolidation of LTF. Adapted from Zhang et al. (2021). Blue denotes the ERK/RSK/p38 MAPK feedback loop, green denotes the PKA/NT/Trk feedback loop. Red denotes pathways involving transcription factors CREB1 and CREB2. Dark blue denotes pathways involving TGF-β. Each number represents a signaling pathway. Arrowheads indicate activation; circular ends, repression.

Two feedback loops were proposed in previous studies to induce a sustained increase of PKA activity, one via the NT/TrkB pathway (Fig. 10, pathways 1→3) (Jin et al. 2018), the other via a CREB1/ *Aplysia* ubiquitin C-terminal hydrolase / PKA positive feedback loop (Hegde et al. 1997; Muller and Carew, 1998; Chain et al. 1999; Mohamed et al. 2005). The present study provides evidence for a third feedback loop, involving the CREB1/TGF-β/Smad pathway (Fig. 10, pathways 11→16→18→20) (Fig. 8), with data suggesting a predominant role for this feedback in the late increase in PKA activity.

Combining data in Fig. 5 and from previous studies (Kopec et al. 2015; Zhang et al. 2017, 2021, 2023), we hypothesize that the TrkB and CREB1/TGF-β feedback loops not only sustain PKA activity but also contribute to the dynamics of ERK activity (Fig. 10). An additional, negative feedback loop ERK/RSK/p38 MAPK acts as a constraint on the long-term dynamics of ERK activity (Fig. 10, pathways 5→6→7) (Zhang et al. 2021). It is possible that the second wave of increase in pERK is regulated by a second feedback loop involving TGF-β (Fig. 10, pathways 5→12→16→18→19), temporally distinct from the loop activating R-Smad and PKA.

RSK can be activated by ERK and PKA (Liu et al. 2019; Zhang et al. 2021). However, PKAc and pERK were increased immediately after 5-HT protocols whereas pRSK was not (Fig. 3). RSK activity was significantly increased immediately after treatment only if the RSK inhibitor BI-D1870, or the p38 MAP kinase inhibitor SB 203580, was applied before and during treatment (Fig. 4). Because p38 MAPK is a downstream kinase of RSK (Zhang et al. 2021), these results suggest another negative feedback loop, involving p38 MAPK and RSK, is responsible for the delay in the increase of RSK activity (Fig. 10, pathways 6→8). Although a single pulse of 5-HT induces rapid RSK activation (Zhang et al. 2021), the accumulated increase in pRSK during 5 pulses will induce a substantial increase in p-p38 MAPK, which could activate negative feedback to reduce pRSK immediately after 5-HT treatment.

The results illustrated in Fig. 8 suggest that TGF-β may also induce the second wave of increase in PKAc at 5 h, via a Smad – dependent feedback loop (Fig. 10, pathways 11→16→18→20). Increased PKAc may induce the second wave of increase in pRSK (Fig. 10, pathway 9). In contrast, p-p38 MAPK was increased immediately after 5-HT protocols whereas pRSK was not, indicating that a pathway independent of RSK (e.g., activation by MKK6, Kumar et al. 2021) is responsible for activating p38 MAPK (Fig. 10, pathway 10) (Zhang et al. 2017). Previous data also suggest negative feedback from p38 MAPK to ERK (Zhang et al. 2021) (Fig. 10, pathway 7), however, unlike RSK, pERK increases immediately after 5-HT treatment (Zhang et al. 2023a). Thus, p38 MAPK might be involved in multiple negative feedback loops, with differing strength, that regulate ERK and RSK (Fig. 10, pathways 7 and 8). Likewise, PKA can induce multiple pathways to activate ERK (Fig. 10, pathways 1→2→4) and RSK (Fig. 10, pathway 9) (Zhang et al. 2021). The increase of pERK and pRSK at 1 h was dependent on PKA (Fig. 5), indicating that at 1 h, the strength of activation by PKA overcomes the inhibitory effects of p38 MAPK.

Similar, complex interactions between kinase cascades involved in learning and memory are generally acknowledged across species. For example, the interacting MAPK and PKA cascades in lateral amygdala neurons are required for consolidation of fear memory (Johansen et al. 2011). The biological significance of feedback loops affecting kinase activation has been widely discussed, for invertebrate and vertebrate systems (Aslam et al. 2009; Angel et al. 2011; Rosenwasser and Turek 2015; Smolen et al. 2019; Hastings et al. 2020; Sump and Brickner 2022; Zhao et al. 2023; Bruce et al. 2024). Strong positive feedback loops have been proposed to induce bistable switches after a transient stimulus, which can permanently switch kinase activities between steady states (Bhalla and Iyengar 1999; Lisman and Zhabotinsky 2001; Markevich et al. 2004; Song et al. 2007). Positive feedback loops can also be essential for converting brief stimuli into long-lasting responses (Smolen et al. 2006). Positive feedback is, therefore, important for converting brief stimuli into long-term synaptic potentiation and memory. In contrast, negative feedback loops can constrain kinase activation and the extent of synaptic plasticity. Crosstalk between positive and negative feedback might help maintain a stable but re-activatable memory trace. In addition, functionally overlapped feedback loops are likely to increase the resistance of consolidated memory to environmental perturbations. If one loop or pathway was suppressed, other pathways might take compensatory roles to ensure memory was consolidated and maintained.

### Functional significance of the dynamic signatures of these cognitive kinases

One of the striking features of the results is the elevation of pRSK 24 h after the E protocol. Although the functional significance of this late wave is yet to be elaborated, it may be critical for mediating the extended LTF produced by the E protocol (Zhang et al. 2011). All three protocols were associated with a gap between the first and second waves. This gap may correspond to a transition between intermediate-term facilitation (ITF) and LTF (Byrne and Hawkins 2015).

The dynamic profiles of kinases may also set the times of eligibility for a second learning trial to further consolidate a memory. The existence of cellular and population clocks for biological timekeeping is commonly accepted (Buonomano and Laje 2010; Gliech and Holland 2020). Multiple waves of kinase activation occur within 24 h in response to 5-HT. After 18 h, we speculate that the inhibitory negative feedback loops lead to inactivation of the kinases, so that LTF does not persist at late times after the Standard protocol (e.g., 48 h, Zhang et al. 2011). The Enhanced protocol seems to modify the timer such that increases of ERK and RSK activities persist to at least 24 h, which may be responsible for maintaining LTF for 5 days (Zhang et al. 2011).

The effectiveness of spaced learning, with multiple repetitions of a stimulus, depends on the time intervals between repetitions (reviewed in Smolen et al. 2016). The feedback loops discussed above are likely important for maintaining an “eligibility trace”, consisting in part of individual or combined activation of the kinases studied here, or of their downstream transcription factors such as CREB1/2. This trace would enable subsequent stimulus repetitions to reinforce synaptic plasticity and memory. For example, two or more blocks of the Standard protocol, spaced 24 h apart, prolong LTF and LTM up to 7 days (Hu et al. 2011, 2015), suggesting priming of the second block by a trace 24 h after the first block. The rich dynamics of the kinases also predict that the effectiveness of a second block of trials might be very dependent on the timing of the second block. For example, a second block of 5-HT stimuli at 18 h after the first might not be effective, because it would come at a time when p38 MAPK is active and inhibiting transcription *via* activation of CREB2 (Guan et al. 2002, 2003; Zhang et al. 2017). Further investigation will be needed to test these hypotheses.

## Materials and Methods

### Neuronal cultures

Invertebrate hermaphrodite *Aplysia californica* (NIH *Aplysia* resource facility, University of Miami, Miami, FL) were maintained in circulating artificial seawater (ASW) at 15°C. Sensory neurons (SNs) were isolated from the ventral-caudal cluster of the pleural ganglion from 60-100 grams *Aplysia* following standard procedures (Zhang et al. 2017, 2021; Liu et al. 2020). Falcon petri dishes of 50 mm x 9 mm were used for cultures. Each dish was plated with 10-15 SNs.

Before experiments, SN cultures were maintained in growth medium (50% adjusted L15 with 1.2 mg/ml of L-Glutamine and 50% hemolymph) for 5-6 days at an 18°C incubator (Thermo Scientific Forma 3900), in dark. Because not all the SNs plated in dishes would grow, we checked the dishes before experiments. Dishes were excluded if they contained less than 5 viable SNs. At least two hours before the treatments, the growth medium was replaced with a solution of 50% L15 and 50% ASW (L15-ASW, including 450 mM NaCl, 10 mM KCl, 11 mM CaCl2, 29 mM MgCl2, 10 mM HEPES at pH 7.6).

### Immunofluorescence analysis

Immunofluorescence procedures for SNs followed those of Zhang et al. (2017, 2021). After the treatments, cells were fixed in a solution of 4% paraformaldehyde in PBS containing 20% sucrose, followed by PBS wash. Fixed cells were then blocked for 30 min at room temperature in a solution of Superblock buffer (Pierce) containing 0.2% Triton X-100 and 3% normal goat serum. Cells were subsequently incubated with primary antibodies overnight at 4° C. Primary antibodies used in this study were: anti-cAMP protein kinase catalytic subunit (PKAc) (anti-PKAc, Abcam, Cat # ab76238, RRID: AB_1523259, 1:400 dilution), anti-phosphorylated ERK (anti-pERK, Cell Signaling, Cat # 4370, RRID: AB_2315112, 1:400), anti-phosphorylated RSK (anti-pRSK, Cell Signaling, Cat # 9346, RRID: AB_330795, 1:400), and anti-phosphorylated p38 MAPK (anti-p-p38 MAPK, Cell Signaling, Cat # 4511, RRID: AB_2139682, 1:400) rabbit antibody. The specificity of these antibodies has been validated in previous studies (Fioravante et al. 2005; Liu et al. 2019; Zhang et al. 2017, 2021, 2023).

After PBS wash of primary antibodies, cells were incubated with secondary antibody (goat anti-rabbit secondary antibody conjugated to Rhodamine Red, Jackson ImmunoResearch Lab, Cat # 111-295-144, RRID: AB_2338028, 1:200) for 1 h at room temperature. Cells were then mounted using Mowiol 4-88 (SigmaAldrich) and dried at room temperature, protected from light, before images were taken. Images of cells were obtained with a Zeiss LSM800 confocal microscope using a 63x oil-immersion lens. A laser with excitation wavelength of 561 nm was used for images acquisition. Gain and offset were adjusted to prevent the saturation of fluorescent signals. A z-series of optical sections through the cell body (0.1 µm increments) were taken, and the single section through the middle of the soma was used for analysis of mean fluorescence intensity. Each image had a resolution of 1024 x 1024 pixels, 8 bits per pixel. In experiments where cultures were paired, or grouped, image acquisition settings were kept identical across samples. The section through the middle of the soma was used for quantification, as in our previous studies and others (Liu et al. 2008, 2011; Bougie et al. 2009; Zhang et al. 2011, 2023; Miranda et al. 2023), because *Aplysia* SNs vary in cell size, and standardizing the region of analysis can minimize the effect of this variation.

ImageJ-win64 software (NIH) was used for measuring the mean fluorescence intensity in the soma acquired in optical sections. All the cells on each coverslip were analyzed and averaged. The number of samples (n) reported in Results indicates numbers of dishes quantified.

### Experimental design

Dishes of SNs cultured from the same animal were paired for all the 5-HT treatments. One dish received L15-ASW as vehicle control (Veh). The other received the same solution with the addition of 5-HT. Three 5-HT protocols were applied to SNs. In each protocol, one pulse of 5-HT consisted of a 5-min bath application of 50 µM 5-HT (Sigma). At the end of the pulse, 5-HT was washed with 10 ml of L15/ASW.

In experiments to measure time courses of phosphorylated RSK (pRSK), phosphorylated p38 MAPK (p-p38 MAPK), or PKA catalytic subunits (PKAc) after 5-HT protocols, one of each pair of dishes was either fixed for immunofluorescence immediately after the end of treatment, or incubated in L15/ASW after wash off of 5-HT until fixation at specified times (1 h, 2 h, 5 h, 18 h and 24 h). The other dish served as a time-matched Veh control. For each pair of dishes measured at the same time point, the averaged level of immunofluorescence intensity from the dish receiving 5-HT was compared to the averaged value from the Veh control. Technical challenges limited our ability to visualize the activation of two or more kinases simultaneously within the same neuron.

### Pharmacological treatment

To ensure sufficient time for inhibitors to penetrate the cells, inhibitor applications began at least 0.5 h before and continued until the end of 5-HT treatment. For post-treatment time points, inhibitors were applied 1 h prior to the specified times at which cells were fixed for immunofluorescence, and remained until fixation. To examine negative feedback effects on pRSK immediately after 5-HT treatment, 3 μM p38 MAPK inhibitor SB203580 (SB) (Sigma) or 4 μM RSK inhibitor BI-D1870 (BID) (Santa Cruz) was applied 0.5 h before 5-HT treatment and continued during treatment. These concentrations were used in previous studies to block the effects of RSK or p38 MAPK (Zhang et al. 2017; Liu et al. 2020, 2022).

To examine the effects of PKA activity on pRSK and pERK at 1 h after 5-HT treatment, 10 μM KT5720 (Sigma) was applied to SN cultures 0.5 h before and during the treatment. To examine the effects of PKA activity on pRSK at 5 h after 5-HT treatment, 10 μM KT5720 was applied to SN cultures 1 h prior to fixation. At this concentration, KT5720 inhibits PKA activity in *Aplysia* without affecting basal synaptic strength (Zhang et al. 2021).

To examine the effects of the MEK-ERK pathway on the activity of RSK at 5 h after 5-HT treatment, the MEK1/2 inhibitor U0126 (Cell Signaling) (20 μM) was applied to SN cultures 1 h prior to fixation. At this concentration, U0126 blocks ERK phosphorylation after 5-HT treatment in *Aplysia* (Sharma et al. 2003; Zhang et al. 2017).

To examine the effects of TrkB on pRSK at 5 h after 5-HT treatment,10 μg/ml of a TrkB antagonist, TrkB Fc chimera (via binding to and neutralizing TrkB ligands, preventing ligand-mediated signaling) (TrkB Fc) (R&D Systems), was applied to SN cultures 1 h prior to fixation. Previously, at this concentration, TrkB Fc inhibited the increase of pERK in isolated *Aplysia* SNs 18 h after the Standard and two-pulse protocols, and 24 h after the Enhanced protocol, without affecting basal activity (Zhang et al. 2023a).

To examine the effects of TGF-β on pRSK or PKAc at 5 h after 5-HT treatment, 5 μg/ml of TGF-β RII Fc chimera (TGF-β RII Fc) (R&D Systems), was applied to SN cultures 1 h prior to fixation. At this concentration, TGF-β RII Fc inhibited the increase of pERK in isolated *Aplysia* SNs 18 h after the Standard and two-pulse protocols, and 24 h after the Enhanced protocol, without affecting basal activity (Zhang et al. 2023a).

To examine the effects of Receptor-regulated mothers against decapentaplegic homolog (R-Smad) protein in *Aplysia*, the analog of Smad3 in mammalian cells, on PKAc at 5 h after 5-HT treatment, 10 μM of Smad3 inhibitor SIS3 (Sigma), was applied to SN cultures 1 h prior to fixation. At this concentration, SIS3 efficiently inhibited the effects of Smad3 in vertebrate and invertebrate cells (Qureshi et al. 2008; Boudreau et al. 2012; Wu et al. 2018; Li et al. 2020).

For each inhibitor experiment described above, four dishes of SNs from the same animal were used. Each dish was given a different treatment, either: 1) 50 μM 5-HT alone; 2) inhibitor alone; 3) 5-HT + inhibitor; or 4) Veh alone. The number of samples (n) reported in Results indicates the number of animals. At least five animals were used in each experiment. All treatments and measures were performed at daytime, at room temperature, and compared to time matched controls, therefore the identified changes in kinase activities were not likely due to a circadian effect.

### Statistical analyses

SigmaPlot version 11 (Systat Software) was used for statistical analyses. Before applying statistical tests, Shapiro-Wilk Normality and Equal Variance tests were performed. In the experiments to compare kinase activity between paired Veh and 5-HT treatment groups at different times, if data passed normality and equal variance tests at all time points, a paired t-test with Bonferroni corrections was used for comparison between paired groups. Otherwise, a Wilcoxon Signed Rank Test (WSRT) with Bonferroni corrections was used. The p value adjusted by Bonferroni corrections was the p value obtained after paired t-test multiplied by the number of groups. If the adjusted p-value exceeded 1, p was reported as 1.

When multiple comparisons of kinases at 18 h or 24 h were performed between groups treated with three different 5-HT protocols, one-way ANOVA and the post hoc Student-Newman– Keuls (SNK) tests. Kruskal-Wallis one way analysis of variance on ranks was used if data displayed a non-normal distribution.

When multiple comparisons between groups treated with 5-HT and inhibitors were performed, if data passed normality and equal variance tests, one-way repeated measures (RM) ANOVA and the post hoc SNK method were used on raw data. If data displayed a non-normal distribution, Friedman repeated measures analysis of variance on ranks and the post hoc SNK method were used.

In all immunofluorescence experiments, the experimenter was blind to the identity of treatments given to each dish of cells. A p value less than 0.05 was considered to represent statistical significance. The data from this article will be shared on reasonable request to the corresponding author.

## Acknowledgements

We thank E. Kartikaningrum for preparing the cultures, C. Neveu for assistance with the illustrations, and R. Calvo for comments on the manuscript. This research was supported by NIH grants NS019895 and NS102490.

